# Distinctive associations between plasma p-tau181 levels and hippocampal subfield volume across the Alzheimer’s disease continuum

**DOI:** 10.1101/2025.01.27.635113

**Authors:** Adea Rich, Hwamee Oh, Alzheimer’s Disease Neuroimaging Initiative

**Affiliations:** Brown University School of Public Health; Departments of Psychiatry and Human Behavior and Cognitive and Psychological Sciences, Brown University; Carney Institute for Brain Science, Brown University

## Abstract

**Background:** Plasma p-tau181 is a promising diagnostic marker of Alzheimer’s disease (AD) pathology, reflecting amyloid accumulation, tau deposition, and downstream neurodegeneration that leads to cognitive impairment. However, the specificity of plasma p-tau181 to AD-related tau pathology remains unclear.

**Objective:** To assess whether plasma p-tau181 is differentially associated with volumetric changes in distinct hippocampal subfields and whether they mediate the relationship between plasma p-tau181 and cognition across the AD continuum.

**Methods:** 213 participants with normal cognition (N=57), mild cognitive impairment (N=109), and AD (N=47) from the Alzheimer’s Disease Neuroimaging Initiative (ADNI) were included for cross-sectional analyses of hippocampal subfield volume that was quantified using the Automatic Segmentation of Hippocampal Subfields (ASHS) software. A subset (n=89) was evaluated for one-year longitudinal changes in hippocampal subfield volume.

**Results:** Higher plasma p-tau181 levels (pg/mL) were associated with decreased volumes in the CA1 and dentate gyrus, bilaterally, and right entorhinal cortex (*ps* < 0.05). Additionally, volumes of these subfields partially mediated the relationship between plasma p-tau181 and ADNI memory and executive function composite scores. Baseline plasma p-tau181, however, did not predict longitudinal atrophy of hippocampal subfields across diagnostic groups.

**Conclusions:** Plasma p-tau181 is differentially associated with hippocampal subfields that are closely related to both age- and AD-related neurodegeneration. Elevated plasma p-tau181 levels may reflect tau accumulation, and volumetric changes in CA1 and DG may mediate the detrimental effect of tau pathology on cognition.

## Introduction

Alzheimer’s disease (AD) is a neurodegenerative disorder pathologically characterized by the accumulation of extracellular amyloid-β (Aβ) plaques and intracellular neurofibrillary tangles (NFTs).^1^ Recent advances in amyloid and tau positron emission tomography (PET), as well as Aβ42, phosphorylated-tau (p-tau), and total tau (t-tau) measured in cerebrospinal fluid (CSF), have been shown to accurately detect these neuropathological features.^2–5^ Structural magnetic resonance imaging (MRI) is also a widely validated biomarker for assessing atrophy in the medial temporal lobe, reflecting dendritic and neuronal damage associated with AD.^6–8^ While PET and CSF markers of Aβ and tau have improved diagnostic accuracy for prodromal and dementia stages of AD, there remains a need for non-invasive, cost-effective, and accessible biomarkers that offer high sensitivity and specificity for both clinical and research settings.^9^ Moreover, abnormal levels of Aβ and hyperphosphorylated tau (p-tau) pathologies begin to accumulate 15-20 years before symptom onset.^10,11^ Considering the prolonged preclinical phase of AD, the availability of cost-effective and scalable biomarkers that can assist diagnosis and prognosis for such at-risk populations will be crucial for early diagnosis of the disease.^12^

Several blood-based plasma markers have been developed and assessed to validate their diagnostic and prognostic capabilities.^13–25^ These markers include Aβ40, Aβ42, t-tau, different isoforms of p-tau, such as p-tau181, p-tau217, and p-tau231, and neurofilament light chain (NfL), which show a varying degree of diagnostic and prognostic capabilities along the AD continuum.^26–28^ In particular, the plasma biomarkers p-tau181, p-tau217, and p-tau231 have emerged as promising indicators of Aβ and tau pathologies in AD.^29–31^ Plasma levels of p-tau181, p-tau217, and p-tau231 have been shown to increase with cognitive decline, hippocampal atrophy, and the intensity of Aβ and tau pathologies.^32–36^ Additionally, these biomarkers effectively distinguish between individuals with AD and healthy controls.^37^ All three p-tau measures, however, correlate more strongly with amyloid PET than with tau PET, with p-tau217 demonstrating the strongest correlation, followed by p-tau231 and then p-tau181.^38^ Moreover, compared to p-tau181, plasma p-tau217 and p-tau231 show earlier and stronger associations with Aβ as well as tau pathologies, becoming abnormal in Aβ-positive, and tau-negative (preclinical AD) individuals.^21,33,34,39,40^ Plasma p-tau217 is particularly accurate in diagnosing AD dementia and can distinguish AD from primary age-related tauopathy (PART) and mixed AD pathologies, outperforming plasma p-tau181.^41^ While plasma p-tau231 and p-tau217 show earlier changes in preclinical AD, plasma p-tau181 remains frequently used in clinical trials and is strongly correlated with both amyloid pathology and cognitive decline.^34,42–44^ Although soluble p-tau plasma measures are considered as a promising AD biomarker and are interpreted as reflecting tau pathology, concentrations of p-tau have shown stronger associations with Aβ deposition than tau pathology, especially in cognitively unimpaired individuals.^38,45^ Therefore, it remains unclear to what extent plasma p-tau measures reflect tau neurofibrillary tangles in the brain, especially in the early stage of AD pathologies.

Increased plasma p-tau181 has been linked to hippocampal atrophy,^46–48^ while hippocampal (HC) subfields has been implicated for differential susceptibility to AD.^49–51^ The HC formation includes the entorhinal cortex (ERC), subiculum (SUB), cornu ammonis regions (CA1-3), and dentate gyrus (DG), each structurally diverse with distinct functions and varying disease susceptibility.^52–55^ In AD, atrophy predominantly affects the ERC, CA1, and SUB, while the DG and CA3 are often spared.^50,56,57^ In healthy aging, atrophy typically occurs in DG and CA3.^58^ Examining HC subfield integrity in association with plasma p-tau181 may improve the detection of subtle AD-related structural changes and their cognitive implications,^59,60^ while clarifying how plasma p-tau181 specifically relates to tau pathology that could exert its downstream effects, such as hippocampal atrophy and cognitive decline.^61,62^ Examining the specific relationship between plasma p-tau181 and hippocampal subfield integrity, thus, will help to gain a more comprehensive understanding of the diagnostic and prognostic capabilities of plasma p-tau181.

In this study, we examined how plasma p-tau181 was associated with HC subfield volume across diagnostic groups (cognitively normal, mild cognitive impairment, and AD) and assessed whether baseline plasma p-tau181 could predict HC volumetric changes between baseline and a one-year follow-up. Additionally, we examined whether HC subfield volume mediated the associations between plasma p-tau181 and cognition. We hypothesized that elevated plasma p-tau181 levels would be associated with greater atrophy in CA1, ERC, and SUB across the AD continuum—subfields particularly vulnerable in AD compared to normal aging—and that these subfields would mediate the relationship between plasma p-tau181 and both memory and executive function.

## Methods

### Study participants

Participants were selected from the Alzheimer’s Disease Neuroimaging Initiative (ADNI) database (https://adni.loni.usc.edu/). ADNI, initiated in 2003, is a longitudinal multicenter study collecting clinical, imaging, genetic, and biochemical biomarker data to aid in the understanding of AD.^63^ The project obtained approval from the institutional review boards of all collaborating institutions, and written informed consent was obtained from all participants.

In this study, participants who were classified as cognitively normal (CN) or cognitively impaired, including those with mild cognitive impairment (MCI), and AD dementia, were included. A total of 213 participants with plasma p-tau181 measurements and high-resolution T2-weighted MRI scans available at baseline were selected from the ADNI database. Subsequent HC subfield volume collected around 1 year later was measured for 89 participants. A detailed description of the sample selection can be found in Figure 1. Additional information regarding eligible participants can be found at https://adni.loni.usc.edu/data-samples/adni-data/.

**Figure 1.**
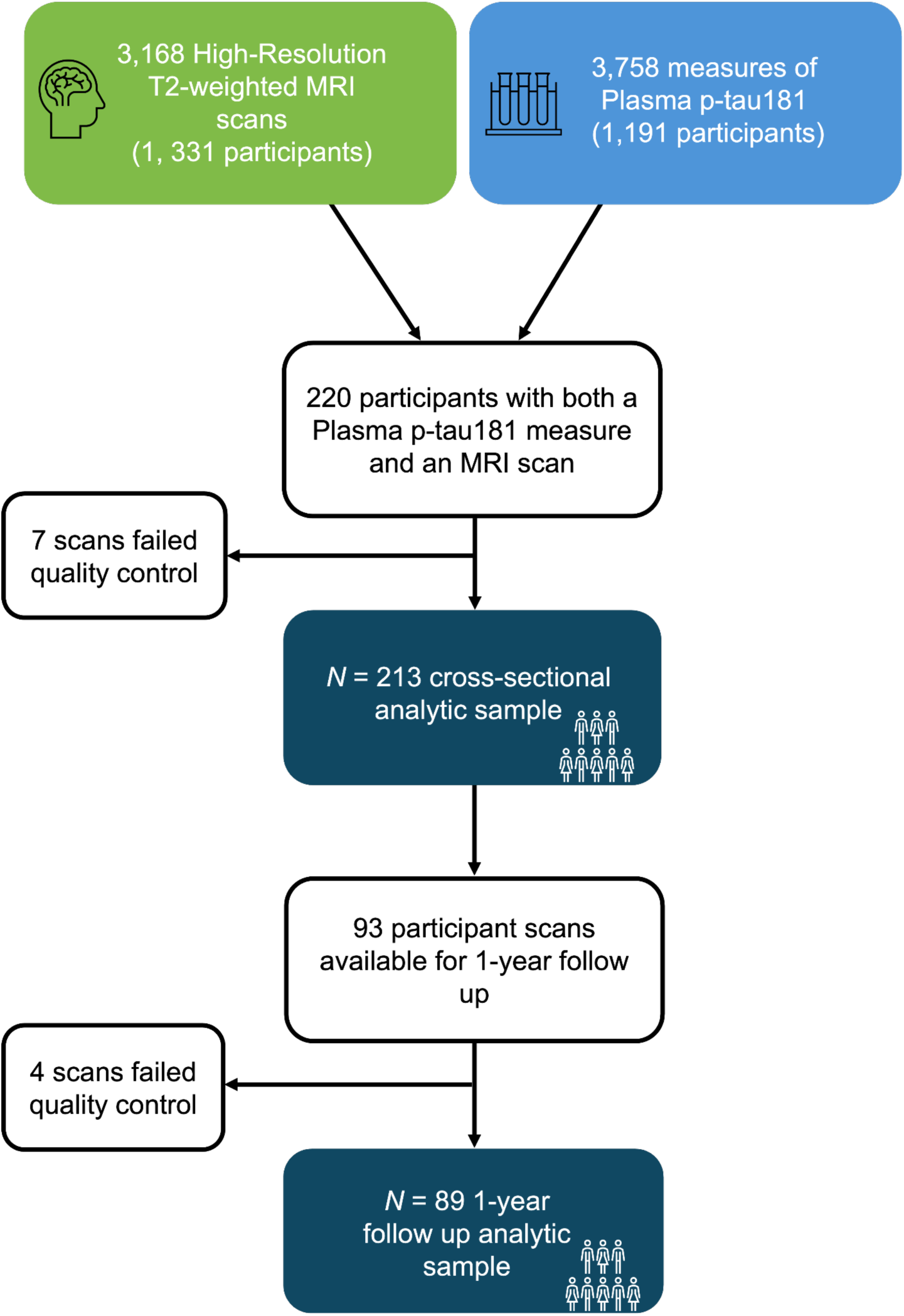
Sample selection from the Alzheimer’s Disease Neuroimaging Initiative (ADNI). 213 participants were selected based on availability of p-tau measures and qualified MRI data cross-sectionally and for the 1-year follow-up.

### Image acquisition

T1-weighted high-resolution MRI scans and T2-weighted hippocampal subfield scans were obtained from various scanners at multiple ADNI sites. Up-to-date information on MRI imaging protocols can be accessed at https://adni.loni.usc.edu/data-samples/adni-data/neuroimaging/mri/.

### Automatic Segmentation of Hippocampal Subfields

We measured HC subfield volumes using the Automatic Segmentation of Hippocampal Subfields (ASHS) software.^64^ The ASHS segments the HC subfields based on a multi-atlas segmentation approach with joint label fusion and bias correction using advanced machine-learning techniques. ASHS is a fully automated framework that includes MRI pre-processing, image segmentation, bias correction, and refinement. It utilizes both high-resolution T1-weighted and T2-weighted images to achieve optimal segmentation and divides the medial temporal lobe into the following subfields: cornu ammonia 1 (CA1), cornu ammonis 2 (CA2), cornu ammonis 3 (CA3), dentate gyrus (DG), subiculum (SUB), entorhinal cortex (ERC), perirhinal cortex of Brodmann area 35 and Brodmann area 36, collateral sulcus, and a miscellaneous part (cysts and cerebrospinal fluid). More information on ASHS and a comparison with the manual approach and FreeSurfer version 6.0 can be found elsewhere.^64^

The volumes of HC subfields used in this analysis (CA1-CA3, DG, ERC, and SUB) were normalized by dividing the raw volumes by the number of slices in which the regions of interest (ROI) appear.^64^ The rate of change for HC subfield volume was calculated by subtracting the adjusted volume at baseline from the adjusted volume at year one and dividing by the time in years between these two measurements. Larger negative values indicate greater atrophy over the (approximately one year) time between data collection.

### Plasma p-tau181

Plasma p-tau181 data were obtained from the ADNI database. Plasma p-tau181 was processed using the clinically validated Single Molecule array (Simoa) HD-X (Quanterix) technique developed at the Clinical Neurochemistry Laboratory, University of Gothenburg, Sweden, as previously described.^48^ This assay utilizes the AT270 monoclonal antibody specific to the threonine-181 phosphorylation site, along with the detection anti-tau mouse monoclonal antibody Tau12, to measure N-terminal to mid-domain forms of p-tau181. The quantification range for this assay, after dilution correction, is between 1.0 and 128.0 pg/mL.

### Cognitive testing

We analyzed ADNI-MEM, a composite memory score based on the Rey Auditory Verbal Learning Task (AVLT), ADAS-COG-13, Mini-Mental State Examination (MMSE), and Logical Memory tasks, as well as ADNI-EF, a composite executive functioning score based on the WAIS-R Digit Symbol Substitution, Digit Span Backwards, Trails A and B, Category Fluency, and Clock Drawing.^65,66^ Higher scores indicate better cognitive performance for both ADNI-MEM and EF.

### Statistical analyses

Statistical analyses were carried out in R version 4.2.2.^67^ Differences among diagnostic groups (CN, MCI, AD) in demographic and clinical characteristics were evaluated using analysis of variance (ANOVA) for numeric variables, the chi-squared test for categorical variables in the cross-sectional sample, and Fisher’s exact test for categorical variables in the 1-year follow-up sample. Differences in HC subfield volumes among diagnostic groups were assessed using analysis of covariance (ANCOVA) and Tukey’s HSD, controlling for age and sex. A linear regression model evaluated the baseline association between plasma p-tau181 concentration and diagnostic groups, adjusting for age and sex. Marginal means were obtained, and Tukey’s HSD assessed pairwise differences, controlling for family-wise error. Another linear regression model examined the association between plasma p-tau181 concentration and HC subfield volume across diagnostic groups, adjusted for age and sex, with adjusted p-values calculated using the Benjamini-Hochberg method to control the false discovery rate (FDR) and account for multiple comparisons. Additionally, a linear regression model assessed baseline plasma p-tau181 concentration and the rate of change in HC subfield volume, adjusted for time (in years), age, and sex. Finally, a mediation analysis determined whether the association between baseline plasma p-tau181 concentration and composite scores for memory and executive function are mediated by HC subfield volume, adjusting for age, sex, and years of education, with bootstrapped 95% confidence intervals. Missing data comprised < 5% for all variables and was handled using complete case analysis.^68^ Statistical significance was defined as *p* < 0.05.

## Results

### Demographic and clinical characteristics

The demographic and clinical characteristics of the participants are summarized in Table 1. Demographic characteristics, including age, education, and sex, did not significantly differ between diagnostic groups in the cross-sectional and 1-year follow-up samples. However, significant differences in clinical characteristics were observed for plasma p-tau181, MMSE, APOE4-carrier status, memory composite score, and executive function composite score among the CN, MCI, and AD groups in both samples (*p* < 0.05). The AD group consisted of more APOE4 carriers and higher plasma p-tau181 levels, and showed lower MMSE, memory composite, and executive function composite scores.

**Table 1.** Demographic and clinical characteristics of subjects.

|  | Cross-sectional sample |  |  |  |  | 1-year follow-up sample |  |  |  |  |
| --- | --- | --- | --- | --- | --- | --- | --- | --- | --- | --- |
|  | Total<br>(N=213) | CN<br>(N=57) | MCI<br>(N=109) | AD<br>(N=47) | p-value | Total<br>(N=89) | CN<br>(N=26) | MCI<br>(N=54) | AD<br>(N=9) | p-value |
| **Sex** |  |  |  |  | 0.32 |  |  |  |  | 0.35 |
| Female | 98<br>(46%) | 26<br>(46%) | 46<br>(42%) | 26<br>(55%) |  | 38<br>(43%) | 8<br>(31%) | 26<br>(48%) | 4<br>(44%) |  |
| Male | 115<br>(54%) | 31<br>(54%) | 63<br>(58%) | 21<br>(45%) |  | 51<br>(57%) | 18<br>(69%) | 28<br>(52%) | 5<br>(56%) |  |
| **Age** | 73.84<br>(7.82) | 74.32<br>(7.43) | 73.44<br>(7.78) | 74.17<br>(8.45) | 0.75 | 73.90<br>(7.95) | 74.50<br>(8.02) | 74.22<br>(7.99) | 70.22<br>(7.38) | 0.34 |
| **Education** | 16.45<br>(2.60) | 16.53<br>(2.56) | 16.56<br>(2.71) | 16.09<br>(2.40) | 0.56 | 16.57<br>(2.71) | 16.38<br>(2.77) | 16.70<br>(2.70) | 16.33<br>(2.83) | 0.85 |
| **APOE4** |  |  |  |  | <0.001 |  |  |  |  | <0.05 |
| Non-carriers | 121<br>(57%) | 43<br>(75%) | 65<br>(60%) | 13<br>(28%) |  | 48<br>(54%) | 18<br>(69%) | 28<br>(52%) | 2<br>(22%) |  |
| Carriers | 92<br>(43%) | 14<br>(25%) | 44<br>(40%) | 34<br>(72%) |  | 41<br>(46%) | 8<br>(31%) | 26<br>(48%) | 7<br>(78%) |  |
| **MMSE** | 26.68<br>(4.01) | 29.00<br>(1.27) | 27.83<br>(2.23) | 21.26<br>(4.50) | <0.001 | 27.40<br>(3.23) | 29.08<br>(0.93) | 27.74<br>(1.82) | 20.56<br>(5.22) | <0.001 |
| **ADNI-MEM** | 0.36<br>(1.14) | 1.24<br>(0.69) | 0.53<br>(0.84) | -1.10<br>(0.70) | <0.001 | 0.51<br>(1.03) | 1.12<br>(0.71) | 0.50<br>(0.87) | -1.23<br>(0.66) | <0.001 |
| **ADNI-EF** | 0.29<br>(1.19) | 0.98<br>(0.85) | 0.48<br>(0.95) | -0.99<br>(1.10) | <0.001 | 0.50<br>(1.11) | 1.00<br>(0.80) | 0.50<br>(1.01) | -0.98<br>(1.16) | <0.001 |
| **P-tau 181**<br>**(pg/mL)** | 18.99<br>(12.99) | 14.58<br>(8.81) | 17.25<br>(10.58) | 28.36<br>(17.29) | <0.001 | 17.60<br>(11.30) | 12.48<br>(8.66) | 18.36<br>(11.39) | 27.85<br>(10.10) | <0.01 |
| **Right CA1 (mm<sup>3</sup>)** | 56.40<br>(11.03) | 62.57<br>(7.78) | 57.38<br>(10.18) | 46.64<br>(9.83) | <0.001 | 56.04<br>(11.09) | 63.12<br>(7.95) | 54.99<br>(10.06) | 41.90<br>(9.54) | <0.001 |
| **Right CA2 (mm<sup>3</sup>)** | 1.91<br>(0.63) | 2.23<br>(0.44) | 1.85<br>(0.65) | 1.66<br>(0.65) | <0.001 | 1.92<br>(0.63) | 2.17<br>(0.67) | 1.87<br>(0.57) | 1.39<br>(0.58) | <0.01 |
| **Right CA3 (mm<sup>3</sup>)** | 8.78<br>(4.01) | 8.31<br>(2.47) | 9.47<br>(4.57) | 7.75<br>(3.90) | <0.05 | 8.72<br>(4.49) | 9.09<br>(4.95) | 8.54<br>(4.01) | 8.73<br>(5.75) | 0.88 |
| **Right DG (mm<sup>3</sup>)** | 42.56<br>(10.32) | 48.18<br>(8.49) | 43.15<br>(9.55) | 34.38<br>(8.97) | <0.001 | 42.37<br>(11.44) | 47.40<br>(8.40) | 41.91<br>(11.31) | 30.59<br>(11.48) | <0.001 |
| **Right ERC**<br>**(mm<sup>3</sup>)** | 35.56<br>(8.87) | 40.05<br>(7.56) | 35.64<br>(8.56) | 29.92<br>(7.98) | <0.001 | 35.73<br>(8.57) | 40.52<br>(6.13) | 35.20<br>(8.17) | 25.06<br>(6.57) | <0.001 |
| **Right SUB**<br>**(mm<sup>3</sup>)** | 21.37<br>(4.19) | 23.30<br>(2.99) | 22.02<br>(3.73) | 17.52<br>(4.01) | <0.001 | 21.15<br>(3.93) | 23.19<br>(3.30) | 20.94<br>(3.66) | 16.57<br>(3.12) | <0.001 |
| **Left CA1 (mm<sup>3</sup>)** | 54.16<br>(10.21) | 59.97<br>(6.98) | 55.05<br>(9.30) | 45.08<br>(9.49) | <0.001 | 54.76<br>(10.52) | 61.57<br>(6.42) | 53.83<br>(9.90) | 40.64<br>(7.53) | <0.001 |
| **Left CA2 (mm<sup>3</sup>)** | 2.59<br>(0.56) | 2.70<br>(0.40) | 2.55<br>(0.60) | 2.54<br>(0.63) | 0.23 | 2.54<br>(0.51) | 2.68<br>(0.44) | 2.54<br>(0.55) | 2.23<br>(0.32) | 0.08 |
| **Left CA3 (mm<sup>3</sup>)** | 8.40<br>(3.55) | 8.31<br>(3.38) | 8.11<br>(3.31) | 9.21<br>(4.19) | 0.20 | 7.64<br>(3.22) | 7.59<br>(2.22) | 7.65<br>(3.46) | 7.71<br>(4.35) | 0.10 |
| **Left DG (mm<sup>3</sup>)** | 40.41<br>(9.93) | 45.33<br>(7.96) | 40.97<br>(9.83) | 33.16<br>(8.12) | <0.001 | 40.86<br>(10.72) | 45.90<br>(8.59) | 40.03<br>(10.97) | 31.26<br>(6.82) | <0.001 |
| **Left ERC (mm<sup>3</sup>)** | 36.69<br>(8.25) | 40.02<br>(6.69) | 37.66<br>(8.28) | 30.40<br>(6.43) | <0.001 | 36.84<br>(8.11) | 41.16<br>(5.65) | 36.14<br>(8.20) | 28.55<br>(6.00) | <0.001 |
| **Left SUB (mm<sup>3</sup>)** | 21.32<br>(4.12) | 23.09<br>(3.40) | 21.80<br>(3.93) | 18.06<br>(3.54) | <0.001 | 21.65<br>(4.33) | 24.32<br>(3.73) | 21.08<br>(4.02) | 17.40<br>(3.23) | <0.001 |

Mean (SD) for continuous variables; n (%) for categorical variables. Abbreviations: CN, Cognitively Normal; MCI, Mild Cognitive Impairment; APOE4, *apolipoprotein E4*; MMSE, Mini-Mental State Examination; ADNI-MEM, Memory composite score; ADNI-EF, Executive Functioning composite score. CA1, Cornu ammonis 1; CA2, Cornu ammonis 2; CA3, Cornu ammonis 3; DG, Dentate gyrus; SUB, Subiculum; ERC, Entorhinal cortex

HC subfield volume for CA1, DG, ERC, and SUB significantly differed among the CN, MCI, and AD groups in both the cross-sectional and 1-year follow-up samples (*p* < 0.001), with the AD group showing lower volumes. Right CA2 also significantly differed among groups in both samples (*p* < 0.01), while left CA2 volume did not show significant differences. Likewise, left CA3 volume did not significantly differ among groups. Right CA3 volume significantly differed among groups in the cross-sectional sample (*p* < 0.05) but did not in the 1-year follow-up sample.

### Hippocampal subfield volume differences among diagnostic groups

Group differences in HC subfield volume among diagnostic groups are presented in Figure 2. Significant differences were observed in CA1, DG, and right ERC subfields among all diagnostic groups (*p* < 0.01), with the MCI and AD groups exhibiting smaller volumes than the CN group. The AD group further demonstrated even smaller volumes than the MCI group. The left ERC and SUB displayed differences between the MCI and AD groups, and between the CN and AD groups (*p* < 0.01), with the AD group showing smaller volumes than the MCI and CN groups. The right CA2 demonstrated significant differences between the CN and MCI groups, and between the CN and AD groups (*p* < 0.01), with the AD group exhibiting smaller volumes than the CN group, and the MCI group exhibiting smaller volumes than the CN group. The right CA3 volume significantly differed between the MCI and AD groups (*p* < 0.05), with the AD group exhibiting lower volumes than the MCI group. The left CA2 and CA3 did not show significant differences among any diagnostic groups.

**Figure 2.**
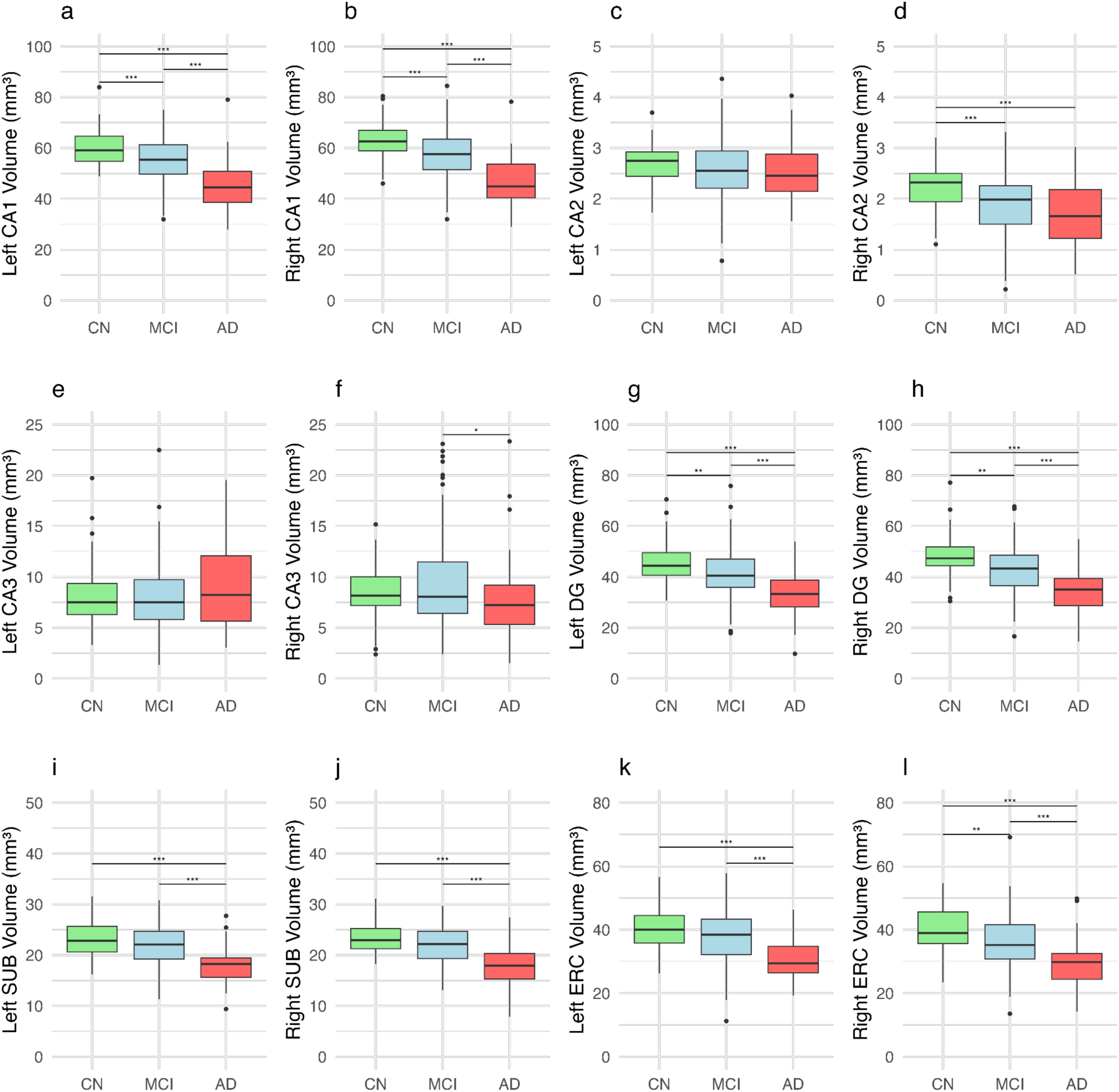
Hippocampal subfield volume differences among diagnostic groups. Panels a-l show the distribution of hippocampal subfield volumes across diagnostic groups. Each box represents the interquartile range (IQR) of the data, with the horizontal line inside the box representing the median. The ‘whiskers’ extend to 1.5 times the IQR above and below the upper and lower quartiles. Abbreviations: CN, Cognitively normal; MCI, Mild cognitive impairment; AD, Alzheimer’s disease; CA1, Cornu ammonis 1; CA2, Cornu ammonis 2; CA3, Cornu ammonis 3; DG, Dentate gyrus; SUB, Subiculum; ERC, Entorhinal cortex. *(*p* < 0.05), **(*p* < 0.01), ***(*p* < 0.001).

### Plasma p-tau181 level differences among diagnostic groups

The estimated marginal means of plasma p-tau181 levels vary among diagnostic groups. Plasma p-tau181 levels are 14.3 pg/mL (95% CI: 11.3 - 17.4) for CN individuals, 17.1 pg/mL (95% CI: 14.8 - 19.3) for individuals with MCI, and 28.5 pg/mL (95% CI: 25.1 - 31.9) for individuals with AD. Pairwise comparisons revealed significant differences between CN and AD (*p* < 0.001) and between MCI and AD (*p* < 0.001), with the AD group having higher plasma p-tau181 concentrations. No significant difference was found between CN and MCI (*p* = 0.34).

### Baseline association between plasma p-tau181 and hippocampal subfield volume across diagnostic groups

Associations between baseline plasma p-tau181 levels and HC subfield volume across diagnostic groups are illustrated in Figure 3. Our findings indicate that higher levels of plasma p-tau181 (pg/mL) are associated with smaller volumes in the left CA1 (*β* = −0.119, *p* = 0.019, Fig. 2a), right CA1 (*β* = −0.154, *p* = 0.005, Fig. 2b), left DG (*β* = −0.120, *p* = 0.016, Fig. 2g), right DG (*β* = −0.146, *p* = 0.005, Fig. 2h), and right ERC (*β* = −0.091, *p* = 0.048, Fig. 2i). In contrast, left CA2 (*β* = 0.002, *p* = 0.491, Fig. 2c), right CA2 (*β* = −0.002, *p* = 0.494, Fig. 2d), left CA3 (*β* = 0.011, *p* = 0.559, Fig. 2e), and right CA3 (*β* = −0.007, *p* = 0.730, Fig. 2f) do not show significant relationships with plasma p-tau181 across diagnostic groups. Similarly, the left SUB (*β* = −0.012, *p* = 0.558, Fig. 2i), right SUB (*β* = −0.032, *p* = 0.144, Fig. 2j), and left ERC (*β* = −0.056, *p* = 0.188, Fig. 2k) do not demonstrate significant relationships with plasma p-tau181 across diagnostic groups. After adjusting for multiple testing, the right CA1 and right DG (*p* < 0.05) remained statistically significant.

**Figure 3.**
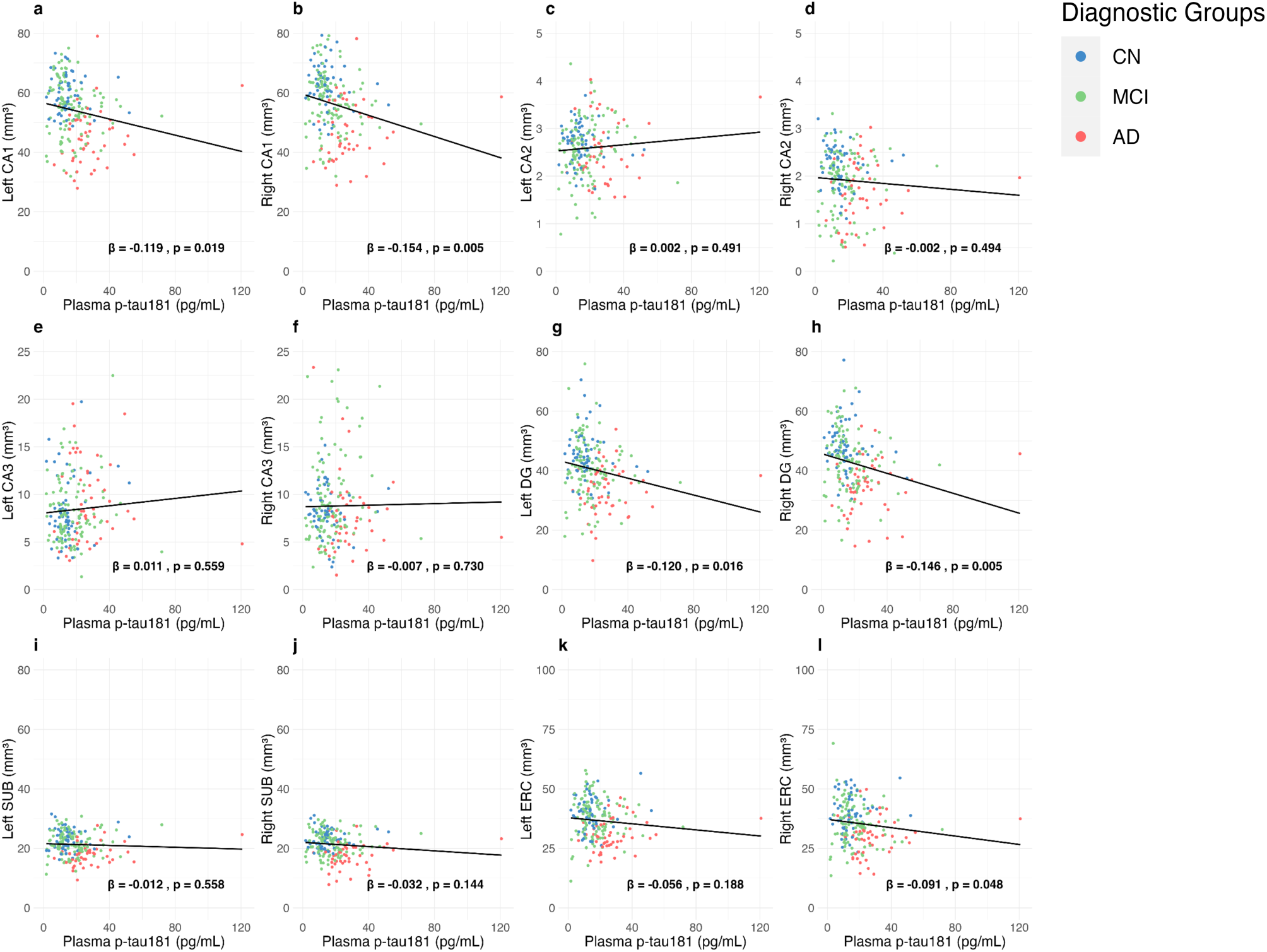
Baseline association between plasma p-tau181 and hippocampal subfield volume across diagnostic groups. *β* coefficients and *p*-values represent the regression model adjusted for age and sex. The scatter plot illustrates the relationship between plasma p-tau181 levels and hippocampal subfield volumes across diagnostic groups without covariates, with each point representing observed data and the line indicating the fitted multiple linear regression model. Abbreviations: CN, cognitively normal; MCI, mild cognitive impairment; AD, Alzheimer’s disease; CA1, Cornu ammonis 1; CA2, Cornu ammonis 2; CA3, Cornu ammonis 3; DG, dentate gyrus; SUB, subiculum; ERC, entorhinal cortex.

### Baseline plasma p-tau181 and hippocampal subfield volume rate of change

There was no association between baseline plasma p-tau181 levels and the rate of change in HC subfield volume from baseline to the 1-year follow-up. Although no subfields demonstrated statistically significant results, several subfields exhibited negative beta values, potentially indicating atrophy over the year: left CA1 (*β* = −0.020, *p* = 0.561), left DG (*β* = −0.100, *p* = 0.109), right DG (*β* = −0.083, *p* = 0.191), left SUB (*β* = −0.066, *p* = 0.128), right SUB (*β* = - 0.061, *p* = 0.170), left CA2 (*β* = −0.002, *p* = 0.726), and right CA2 (*β* = −0.002, *p* = 0.725, Fig. 2d), while other subfields did not exhibit negative beta values: right CA1 (*β* = 0.019, *p* = 0.657), left ERC (*β* = 0.002, *p* = 0.978), right ERC (*β* = 0.036, *p* = 0.462), and right CA3 (*β* = 0.004, *p* = 0.922).

### Mediation analysis

Increased plasma p-tau181 concentration was significantly associated with both lower memory and executive function composite scores (*p* < 0.001). Consequently, a mediation analysis was conducted to assess whether the relationship between plasma p-tau181 and composite scores in memory and executive function is mediated by HC subfield volume. The analysis revealed that CA1, DG, and right ERC volume partially mediate the association between plasma p-tau181 concentrations and both memory and executive function composite scores (Figure 4).

**Figure 4.**
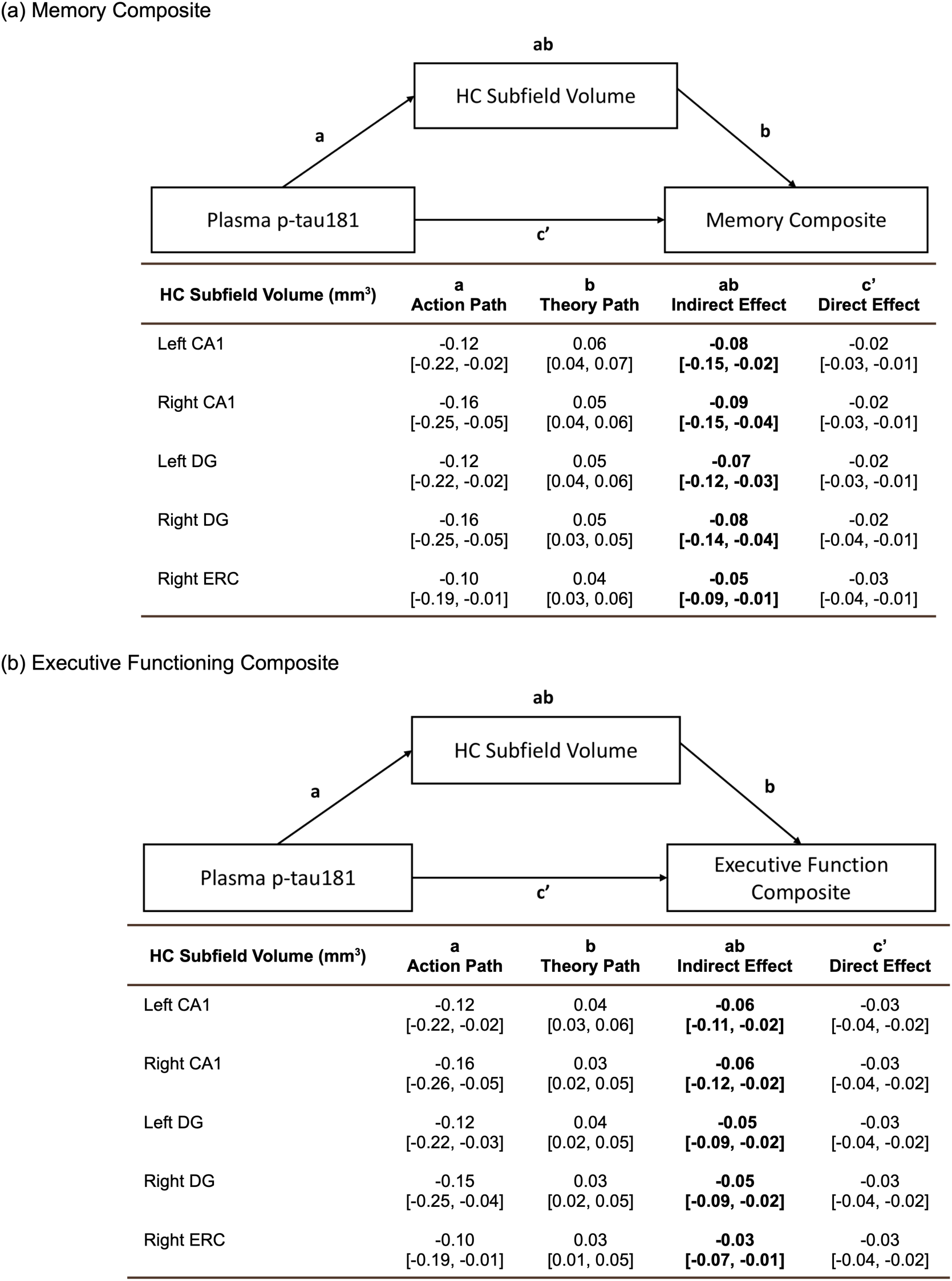
Mediation Analysis. Completely standardized coefficient estimates are presented with bootstrapped 95% confidence intervals in brackets. Mediation analyses revealed a significant indirect effect of hippocampal subfield volume on the relationship between plasma p-tau181 and memory (a), as well as executive functioning (b). While the direct effects of plasma p-tau181 and composite scores were significant, our mediation model demonstrated that elevated plasma p-tau181 concentration was associated with lower hippocampal subfield volumes in CA1, DG, and right ERC, which were linked to lower memory and executive functioning composite scores. Abbreviations: CA1, Cornu ammonis 1; DG, Dentate gyrus; ERC, Entorhinal cortex.

## Discussion

Accumulating evidence indicates that blood-based biomarkers can differentiate diagnostic status across the AD continuum, and they show a high correlation with the severity of AD neuropathologies such as brain Aβ deposition.^69,70^ Among blood-based biomarkers, plasma p-tau181 has been associated with the disease severity of AD with high accuracy, indicating its potential role as a diagnostic biomarker of AD.^71^ Despite its potential utility as a diagnostic biomarker of AD, its relationship with brain tau pathology and a neural mechanism underlying the linkage between plasma p-tau181 and cognitive impairment remain elusive. Based on the previous findings of specific relationships between HC subfields and AD,^52^ we examined how plasma p-tau181 selectively relates to HC subfield volumes at baseline as well as atrophy over one year among older adults across different diagnostic groups (CN, MCI, and AD).

Major findings of the present study include: (1) higher plasma levels of p-tau181 were specifically associated with smaller volume of the CA1, DG, and right ERC cortex cross-sectionally; (2) the volume of these HC subfield regions partially mediated the relationship between plasma p-tau181 and cognitive performance in both memory and executive function; (3) baseline plasma p-tau181 did not predict longitudinal atrophy of the HC subfields. We further replicated that levels of plasma p-tau181 were higher with AD patients compared to CN subjects and that HC subfield volumes known to have differential vulnerabilities to AD pathology, such as the SUB, CA1, and ERC, showed lower volumes in AD groups compared to CN subjects.^71,72^

Our findings of a significant association between higher plasma p-tau181 levels and smaller volumes in CA1, DG, and right ERC subfields are in part consistent with previous research linking AD to volumetric loss in CA1 and ERC.^72,73^ Based on prior evidence suggesting that DG is relatively resistant to AD pathology, we hypothesized that plasma p-tau181 would be associated with reduced CA1 volume, but not DG. In contrast to our prediction, our results did not reveal differential associations between plasma p-tau181 and DG or CA1 volumes across the AD continuum. Interestingly, reductions in both CA1 and DG volumes have been linked to age-related memory decline in older adults, where CA1 integrity has been associated with lower cognitive function scores, and DG with difficulties in pattern separation.^50,57,74,75^ These findings suggest that elevated plasma p-tau181 levels in association with the HC subfields may be more reflective of age-related neurodegenerative processes rather than AD-related tau pathology.

The significant relationship between plasma p-tau181 and DG volume may also point to a potential link with primary age-related tauopathy (PART), which typically involves tau deposition in the DG and CA4 subregions.^76,77^ Conversely, NFTs in DG and CA4 are characteristic of advanced AD, and our findings may indicate neurodegeneration driven by tau pathology in ERC, which disrupts perforant pathway inputs to DG.^78^ Interestingly, while the right ERC demonstrated a significant association with plasma p-tau181, the left ERC did not, and the overall significance observed in ERC was less pronounced compared to DG. Despite plasma p-tau181 being associated with widespread amyloidosis, our results indicate that the relationship between plasma p-tau181 and HC atrophy may closely reflect tau tangles rather than amyloid accumulation. While some studies found a significant association between plasma p-tau levels and temporal meta ROI tau PET levels, these studies included not only nondemented but also demented patients,^38^ potentially indicating that the plasma p-tau levels might have reflected whole-cortex tau tangles in addition to the temporal lobe tau accumulation. Collectively, our results may indicate the link between plasma p-tau181 level and temporal lobe tau accumulation.

Contrary to our hypothesis, we did not observe an association between plasma p-tau181 levels and SUB volume. This finding contradicts existing research that suggests AD pathology and AD blood biomarkers typically correlate with decreased volume in the SUB.^79^

It is plausible a biomarker more specific to AD pathology, such as p-tau217, may be better suited to detect AD-related structural changes. Overall, our findings of the differential association of plasma p-tau181 with HC subfield volume indicate that plasma p-tau181 may be linked to neurodegeneration in both aging and AD, potentially reflecting tau accumulation in HC subfields.^80,81^

Additionally, we found that reduced volumes in CA1, DG, and right ERC partially mediate the relationship between plasma p-tau181 levels and cognitive performance in episodic memory and executive function. Higher plasma p-tau181 levels were associated with lower scores in both cognitive domains, with volumetric decline in these subfields contributing to this association. This is consistent with previous research showing that CA1 and DG mediate the relationship between AD pathology and memory recall and reconsolidation.^51^ Another study found that larger CA1 and CA4-DG volumes predicted better executive function.^82^ In our sample, reduced volumes in CA1 and DG more strongly mediated memory performance than executive function, while both domains were affected. Collectively, our results further support the conclusion that plasma p-tau181 may be associated with broader cognitive impairments,^42^ including both memory performance, such as pattern separation linked to DG atrophy, and executive function.^53,83^ It is important to note that tauopathy in the absence of Aβ pathology, as indicated by increased plasma p-tau181 levels without Aβ, was shown to predict neither cognitive decline nor hippocampal atrophy.^84^ This suggests that Aβ pathology may be a necessary, although not sufficient, component in AD pathophysiology. Despite the absence of Aβ pathology, plasma p-tau181 significantly improved discrimination between Braak I–II and Braak 0 stages among both cognitively unimpaired and impaired individuals, as well as across all older adults combined.^85^ Our findings extend the existing literature by identifying a role of hippocampal subfield volume in the relationship between plasma p-tau181 levels and cognitive impairment and may enhance our understanding of the cognitive difficulties associated with elevated plasma p-tau181 levels.

Baseline plasma p-tau181 levels did not predict longitudinal atrophy in the HC subfields over one year. Previous literature has shown an association between plasma p-tau181 and annual changes in overall HC volume.^32^ However, plasma p-tau181 may not be sensitive enough to detect subtle subfield-specific changes within shorter follow-up periods. In contrast, other AD biomarkers, such as p-tau217 and p-tau231, which exhibit more pronounced increases along the AD continuum, may be better suited for identifying these structural changes. A longer follow-up may be necessary to fully capture the predictive value of plasma p-tau181.

The current study has some limitations. The small sample size may have limited statistical power to detect subfield-specific associations in some HC subfields and restricted our ability to conduct separate analyses within diagnostic groups. Future research utilizing larger and more demographically representative samples could reveal additional relationships between pathology, brain volume, cognitive performance, and plasma biomarkers, particularly with minimal loss to follow-up and sufficient access to track longitudinal changes. Second, although we found a robust relationship between p-tau181 and the HC subfield volumes using the well-validated segmentation tool of the HC subfields, other promising p-tau variants, such as p-tau217 and p-tau231, should also be investigated, which may offer stronger clinical utility due to their ability to detect earlier changes in AD. However, these variants are not currently available in the ADNI dataset, so they could not be assessed as part of this study. Finally, incorporating neuropsychological assessments tailored to the cognitive tasks associated with each HC subfield could further elucidate the cognitive implications of plasma biomarkers.

Our study demonstrates that higher plasma p-tau181 levels are associated with lower volumes in CA1, DG, and right ERC subfields, with these volumes partially mediating the relationship between plasma p-tau181 and cognition. This suggests that plasma p-tau181 levels may indicate tau accumulation, with changes in CA1 and DG mediating the effect of tau pathology on cognition. Identifying which subfields correlate with plasma biomarkers could enhance our understanding of how blood-based plasma biomarkers reflect brain AD pathology and support their diagnostic capabilities. While further research is warranted to clarify these associations, our findings contribute to the growing body of literature examining the relationships between biomarkers, brain structure, and cognitive impairment.

## Author Contributions

**Adea Rich** (Conceptualization; Methodology; Validation; Formal analysis; Writing - Original Draft; Writing - Review & Editing; Visualization), **Hwamee Oh** (Conceptualization; Methodology; Validation; Writing - Original Draft; Writing - Review & Editing; Supervision; Funding acquisition; Project administration)

## Acknowledgments

The authors thank Drs. Zachary Kunicki and Jenna Blujus for their inputs for statistical modeling.

Data collection and sharing for the Alzheimer’s Disease Neuroimaging Initiative (ADNI) is funded by the National Institute on Aging (National Institutes of Health Grant U19AG024904). The grantee organization is the Northern California Institute for Research and Education. In the past, ADNI has also received funding from the National Institute of Biomedical Imaging and Bioengineering, the Canadian Institutes of Health Research, and private sector contributions through the Foundation for the National Institutes of Health (FNIH) including generous contributions from the following: AbbVie, Alzheimer’s Association; Alzheimer’s Drug Discovery Foundation; Araclon Biotech; BioClinica, Inc.; Biogen; BristolMyers Squibb Company; CereSpir, Inc.; Cogstate; Eisai Inc.; Elan Pharmaceuticals, Inc.; Eli Lilly and Company; EuroImmun; F. Hoffmann-La Roche Ltd and its affiliated company Genentech, Inc.; Fujirebio; GE Healthcare; IXICO Ltd.; Janssen Alzheimer Immunotherapy Research & Development, LLC.; Johnson & Johnson Pharmaceutical Research & Development LLC.; Lumosity; Lundbeck; Merck & Co., Inc.; Meso Scale Diagnostics, LLC.; NeuroRx Research; Neurotrack Technologies; Novartis Pharmaceuticals Corporation; Pfizer Inc.; Piramal Imaging; Servier; Takeda Pharmaceutical Company; and Transition Therapeutics.

## Consent to participate

This study was done using publicly obtainable secondary data. Informed consent was obtained through the Alzheimer’s Disease Neuroimaging Initiative.

## Data Availability

The data supporting the findings of this study are openly available in [Ptau181_HC] at [https://github.com/adearich/Ptau181_HC]. These data were derived from the following resources available in the public domain: [https://adni.loni.usc.edu/].

